# A biologically inspired architecture with switching units can learn to generalize across backgrounds

**DOI:** 10.1101/2021.11.08.467807

**Authors:** Doris Voina, Eric Shea-Brown, Stefan Mihalas

## Abstract

Humans and other animals navigate different landscapes and environments with ease, a feat that requires the brain’s ability to rapidly and accurately adapt to different visual domains, generalizing across contexts/backgrounds. Despite recent progress in deep learning applied to classification and detection in the presence of multiple confounds including contextual ones [25, 30], there remain important challenges to address regarding how networks can perform context-dependent computations and how contextually-invariant visual concepts are formed. For instance, recent studies have shown artificial networks that repeatedly misclassified familiar objects set on new backgrounds, e.g. incorrectly labelling known animals when they appeared in a different setting [3]. Here, we show how a bio-inspired network motif can explicitly address this issue. We do this using a novel dataset which can be used as a benchmark for future studies probing invariance to backgrounds. The dataset consists of MNIST digits of varying transparency, set on one of two backgrounds with different statistics: a Gaussian noise or a more naturalistic background from the CIFAR-10 dataset. We use this dataset to learn digit classification when contexts are shown sequentially, and find that both shallow and deep networks have sharply decreased performance when returning to the first background after experience learning the second – the *catastrophic forgetting* phenomenon in continual learning. To overcome this, we propose an architecture with additional “ switching” units that are activated in the presence of a new background. We find that the switching network can learn the new context even with very few switching units, while maintaining the performance in the previous context – but that they must be *recurrently* connected to network layers. When the task is difficult due to high transparency, the switching network trained on both contexts outperforms networks without switching trained on only one context. The switching mechanism leads to sparser activation patterns, and we provide intuition for why this helps to solve the task. We compare our architecture with other prominent learning methods, and find that elastic weight consolidation is not successful in our setting, while progressive nets are more complex but less effective. Our study therefore shows how a bio-inspired architectural motif can contribute to task generalization across context.

## 1 Introduction

The ability to adapt to changes of context while preserving relevant information that is contextually-invariant is one of the traits that make biological brains so effective and robust in complex, natural environments. In the visual domain, the background details or statistics can become irrelevant as agents learn to distinguish visual concepts and act upon them across different settings. For example, humans are particularly adept at generalization across context and will react similarly when seeing a familiar face during a remote video call or during an interaction in person. Translating this capacity for contextual adaptation to artificial intelligent agents is essential if, for example, we are to upgrade current visual learning algorithms to generalize across new environments and abstract visual concepts [3]. One difficulty in this setting is that most current deep feedforward architectures learn the background and foreground in tandem, without abstracting objects of interest from their surround. As we will show, simply relying on network depth is insufficient. Instead, novel architectures supporting context-dependent computations are required.

Another obstacle in achieving general, context dependent processing is that datasets used to train current state of the art neural networks have been shown to be biased [56, 1, 12, 8], with object class correlating with background, or the background statistics being constrained to belong to a particular distribution. For example, certain datasets may predominantly contain street or nature scenes, or have a preferred viewing angle ([54]); in the case of ImageNet, the dataset predominantly contains centered objects with limited background clutter, while the PascalVOC dataset depicts more complex scenes with multiple objects and a significant amount of background clutter [39]. Such biases prevent even the most sophisticated deep architectures from generalizing across contexts, so that when the network is tested on a dataset with different background statistics the accuracy can be significantly affected [2]. Few studies [3, 2] so far have disentangled different dataset biases to separately address how tasks like object classification are affected by variations of context.

To address these challenges systematically, we first construct a simple dataset where context is clearly defined and without other complicating biases. The dataset consists of MNIST digits set on either Gaussian noisy backgrounds, or more naturalistic backgrounds of images from the CIFAR-10 dataset (Figure 1B,D). To parametrically vary the difficulty of this task, we control the transparency of the MNIST digits. We use this dataset in a biologically realistic setting where agents learn to classify digits set on different backgrounds, but are exposed to these contexts sequentially and without having access to previous data points. This is the continual learning framework, where a major challenge is that old tasks are forgotten when network weights are overwritten to solve new tasks, a phenomenon called *catastrophic forgetting*. We show that both deep and shallow networks trained to classify MNIST digits set on the two contexts fail to sequentially learn without catastrophic forgetting. We refer to this learning task with varying context as *sequential context switching*.

**Figure 1:**
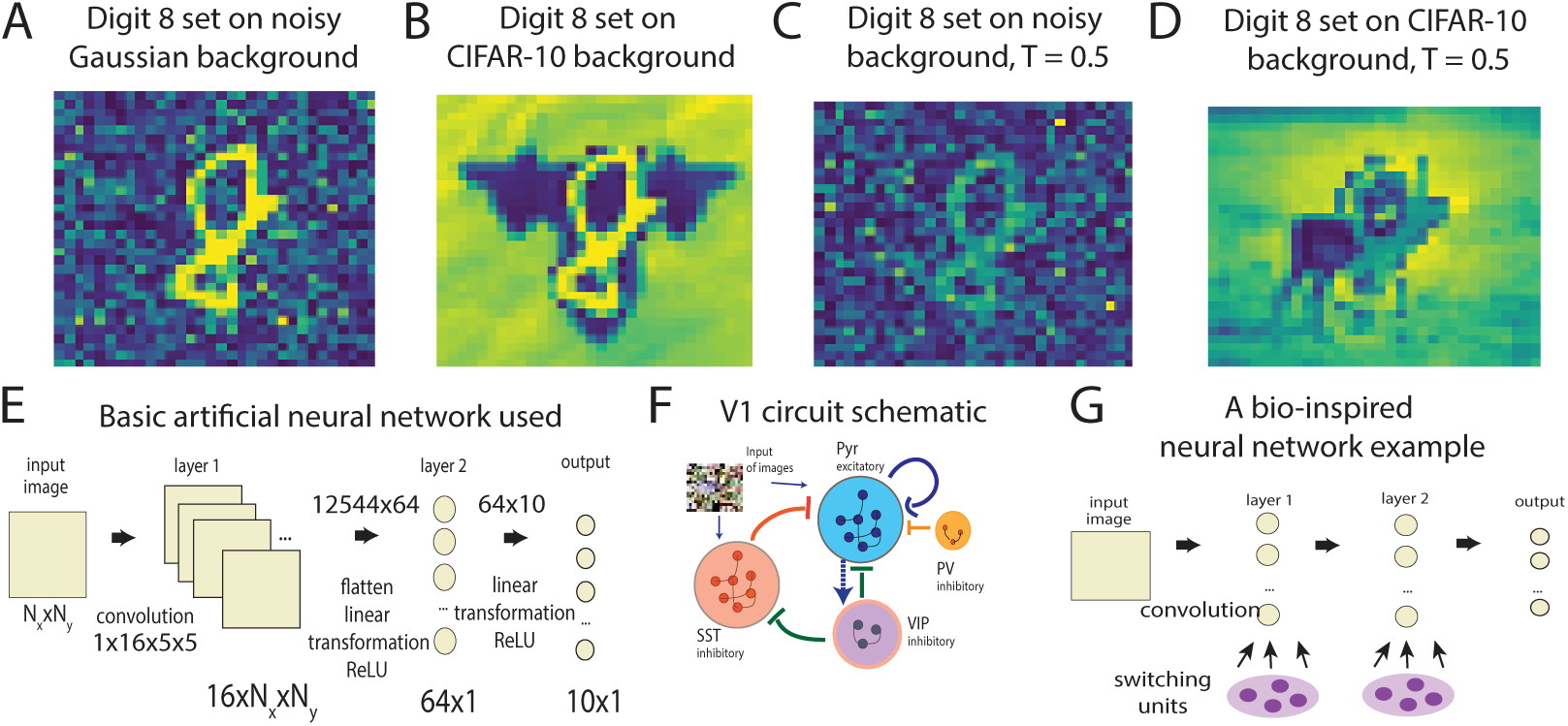
A). Digit “ 8” set on a noisy Gaussian background. B). Digit “ 8” set on a background from the CIFAR-10 dataset. C). Digit “ 8” set on a noisy background with transparency *T* = 0.5. D). Digit “ 8” set on a CIFAR-10 background with transparency *T* = 0.5. E). Basic artificial neural network architecture without switching units used for the toy examples presented below. F). Schematic of the V1 circuit [65] G). A bio-inspired neural network with switching units.

We introduce a new network architecture for solving sequential context switching that is inspired by a recently characterized local circuit in the mouse visual cortex. When an animal shifts from stationary to moving behavior, a population of neurons in the primary visual cortex (V1) which belongs to a cell class expressing Vasoactive Intestinal Peptide – the VIP population – increases its activity. It has been hypothesized that this neuron population turns ON like a switch and reconfigures the circuit dynamics to efficiently process the corresponding static vs. moving visual scenes [55, 65]. Following this, we propose a network architecture, the *switching network*, where we add units that are OFF when the network is presented the first context, but turn ON for the second context, similarly to the VIP (Figure 1F,G). We refer to these added units as *switching units*. More precisely, after the network trains on the first context with no switching units active, the weights learned previously remain fixed, and only weights to and from the active switching units are learned when the second context is shown. Due to the design of this network architecture, *catastrophic forgetting is guaranteed not to occur*.

This leads to the three contributions of our paper: (1) We introduce and share a new dataset that highlights the problem of catastrophic forgetting in sequential context switching. (2) We propose a bio-inspired architecture, the *switching network*, that overcomes catastrophic forgetting. (3) We show that the *switching network* has performance competitive or superior to other methods (EWC [24] and ProgNet [46]) in the sequential context switching task.

## 2 Related Work

Transfer learning (TL), domain adaptation (DA), multi-task learning (MTL), and continual learning (CL) are allied fields that have the broad goal of training networks on two or more datasets or tasks, with the common objective of using the knowledge learned from one task (source task) to efficiently learn a different but related task (target task). General strategies [57] applied to this end include trying to correct for the input marginal distribution difference [27, 15, 49, 18, 40, 39] or the conditional distribution difference [11, 53, 63, 7, 16, 32, 59] between the source and target tasks. In this paper, we focus on context switching, when only the features and statistics of the background change. This scenario is most similar to domain adaptation, a particular case of TL (specifically heterogeneous, transductive TL [10, 58]) when the source and target tasks are identical but the source and target features, as well as their distribution are different due to selection bias or distribution mismatch.

A rich literature spanning decades describes *shallow DA* methods aiming to solve the domain shift between the source and target domains by broadly using two strategies [58]: (1) train models on reweighed source samples to reduce the discrepancy [6, 9]; and (2) find a common shared space where the two domain distributions are matched [19, 40]. Current state-of-the-art DA methods use deep networks – *deep DA* – which have reliably outperformed previous strategies. Various approaches are based on the fine-tuning of deep networks, either by aligning the statistical distribution shift [33, 52], or by adjusting the architectures of deep networks [28, 62]. Another popular method is the adversarial deep DA approach, where a domain discriminator that decides whether the data point belongs to the source or the target domains is used to encourage domain confusion through minimizing the distance between the source and target input distributions [31, 4]. Other well-known methods use autoencoders to reconstruct the inputs, focusing on creating a shared representation between the two domains while maintaining the individual characteristics of each [5]. However, these methods do not specifically address generalization across background and are not tested against a common benchmark. Furthermore, to the best of our knowledge, neither of these network architectures are bio-inspired.

Here we focus on the case when the data is observed incrementally as a continuous stream, a scenario matching the CL approach. Unlike learning in TL and MTL, the network in this case does not have access to old data that can be interleaved with the new data, but rather agents continually learn new knowledge across time while retaining previously learned information [41]. There are several categories of computational approaches to CL that permit good generalization and avoid catastrophic forgetting [41]: (i) learning models that regulate levels of plasticity to protect consolidated knowledge through regularization [29, 43, 13]; (ii) adding additional neural resources such as neurons to learn new information [68, 61, 14, 64, 44]; (iii) using complementary learning systems for memory consolidation and experience replay [20, 17, 50, 23]. In the case of regularization methods, a successful strategy has been applied in [24] (for EWC) and in [67] by which more influential parameters are pulled back towards a reference weight with good performance on previous tasks. An example of the second approach, where the network dynamically adds neuronal resources, is Rusu et al.’s ProgNet architecture [46]. This architecture expands through the allocation of new “ column” networks, trained on novel information, and receiving lateral connections from the other columns. We will describe EWC [24] and ProgNet [46] in more detail below.

Direct comparison between these methods is difficult because of a lack of established benchmark datasets and metrics. Many training and evaluation protocols have shifted from MNIST or CIFAR-10 datasets [24, 67, 21, 66], to more challenging datasets (ImageNet [45], MS COCO [30], OpenImages [25]), similar in complexity to realistic settings [24, 29, 38, 34, 35, 22]. However, large datasets also contain substantial contextual biases, in background/context, rotation, viewpoint, etc. and have insufficient controls to ensure networks do not exploit trivial correlations in the data [2, 60, 69, 48, 47]. For instance, in [60] authors find that changing backgrounds in ImageNet significantly decreases average performance, and that choosing backgrounds in an adversarial manner can lead to misclassifying 87.5% of the images. Therefore, an important undertaking is to carefully analyze generalization (or lack thereof) in detection and classification tasks, dissecting the biases neural networks can abuse or misuse (variations in lightning, viewpoints, context/background etc.) [3]. To that end, our work distinctively addresses variations of context for a classification task, when the network has to generalize by ignoring contexts and abstracting MNIST digits.

## 3 Results

### 3.1 A benchmark dataset for generalization across context

To address the problem of sequential context switching, we create a dataset whose only confounding feature is context. We make this dataset publicly available as a benchmark to test generalization across contexts. The dataset consists of MNIST digits with varying degrees of transparency set on either noisy backgrounds, with noise a Gaussian random variable (Figure 1A), or MNIST digits set on a more naturalistic background from the CIFAR-10 dataset (Figure 1B). We refer to the subset of MNIST digits set on noisy backgrounds as “ MNIST+noise” and to the subset of MNIST digits set on CIFAR-10 backgrounds as “ MNIST+ cifar”. The goal is to perform image classification so that neural networks (NN) correctly identify the MNIST digit despite the different backgrounds. A parameter that makes the task more difficult is digit transparency; as we increase transparency, the background interferes with the digit and the identity of the digit becomes more ambiguous (Figure 1C,D).

### 3.2 Catastrophic forgetting in the MNIST+noise and MNIST+cifar datasets

We first choose a basic NN (Figure 1E) that solves the digit classification task with a performance above 90% (≈ 95%) on either MNIST+noise or MNIST+cifar datasets with no digit transparency. The classic MNIST classification task can be performed at high accuracies using even simple linear classifiers, with accuracies as high as 92.4% reported in [26]. However, the dataset we focus on here contains backgrounds and an additional transparency causing the backgrounds to obscure the digits, making this a more difficult task (Figure 2B,E). For all the experiments presented below, we run simulations 10 times for the basic NN and 5 times for VGG-16 and consider the standard deviation which is comparatively small (see Methods, Supplementary Material).

**Figure 2:**
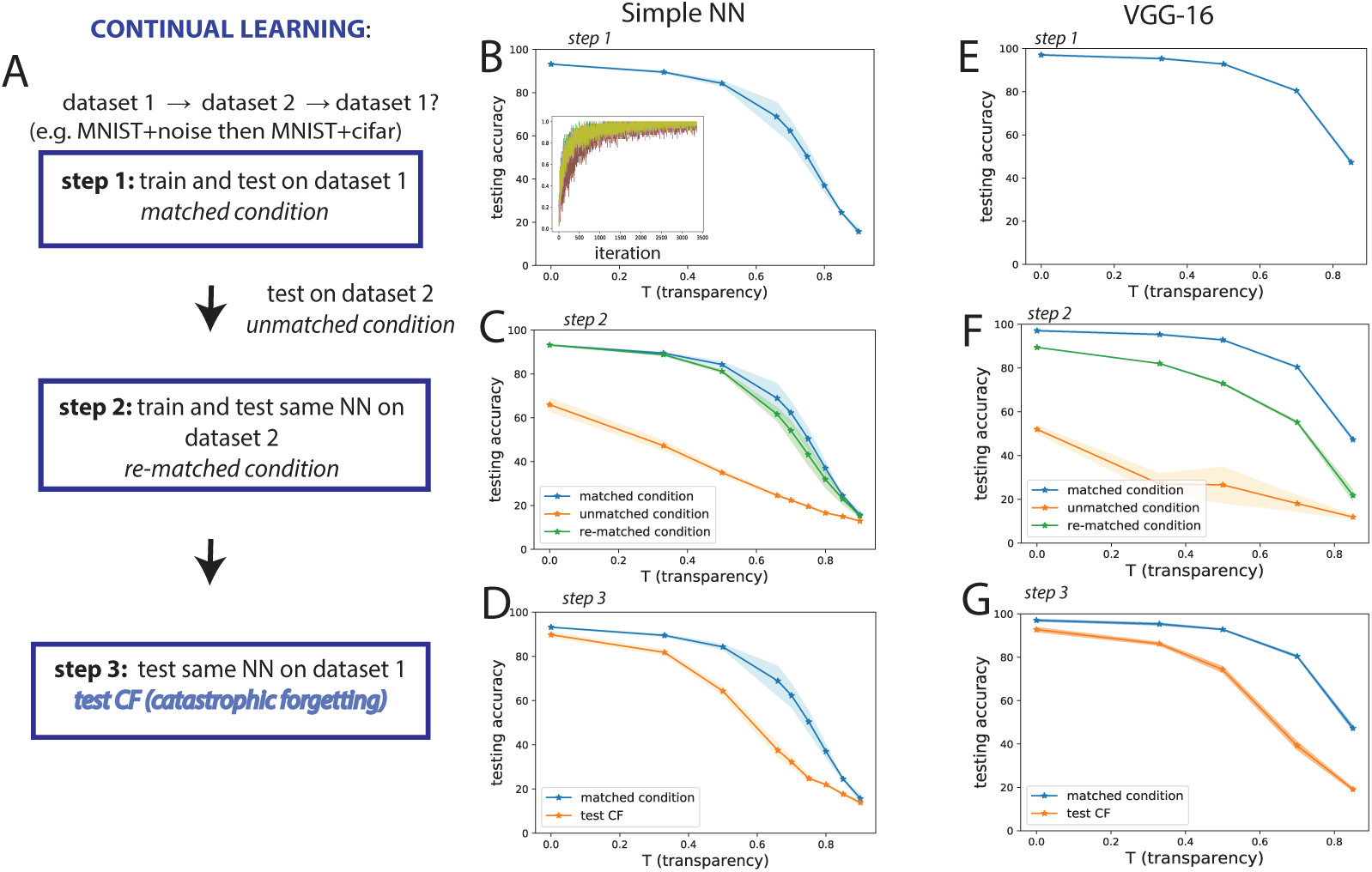
A). Schematic of training and testing for sequential context switching to test catastrophic forgetting. We first train and test on dataset 1 (Step 1), then train and test the same network on dataset 2 (Step 2), then finally test again on dataset 1 to probe forgetting (Step 3). B). Average testing accuracy on the *matched condition* for the MNIST+cifar dataset as transparency *T* increases (inset: convergence during training for the MNIST+cifar dataset). C). The MNIST+cifar dataset is less accurately classified by a network trained on MNIST+noise (*unmatched condition*, orange line) than one trained on MNIST+cifar (*matched condition*, blue line). However, once we re-train the MNIST+noise-trained NN on MNIST+cifar (Step 2), the accuracy approaches that for the *matched condition* (*re-matched condition*, green line). D). The accuracy when re-testing MNIST+cifar on a network that was first trained on MNIST+cifar, then on MNIST+noise, is reduced (*test CF*, orange line) compared to the *matched condition*, therefore we conclude catastrophic forgetting occurs. E)-G) same as B)-D) using the VGG-16 network instead of the basic network.

Turning to the problem of continual learning of context switching, we ask whether this NN learns digit classification on MNIST+noise and MNIST+cifar, sequentially and in either order. To achieve continual learning, at **step 1** we first test dataset 1 on a network trained on dataset 1 – this is the *matched condition*; we then test dataset 2 on a network trained on dataset 1 – this is the *unmatched* condition; at **step 2**, we train the dataset 1-trained NN on dataset 2 and then test on dataset 2 – this is the *re-matched condition*; at **step 3**, we re-test dataset 1 on the NN – this is the test catastrophic forgetting or *test CF condition*, where dataset *i* ∈ {MNIST+noise, MNIST+cifar} (Figure 2A).

Our initial findings are as follows. At step 1, learning digit classification on dataset 1 generates high accuracies (Figure 2B, Suppl. Fig. 1B), in this case up to 97% on the test set (Figure 2B, inset), with reduced accuracy at increased transparency. When testing the NN in the *unmatched condition* using dataset 2, the accuracy is comparatively poor for no training on this context (orange vs blue lines in Figure 2C and Suppl. Fig. 1C). At step 2 for the *re-matched condition*, we obtain a performance (green line, Figure 2C) comparable to the *matched condition*, as if we trained on dataset 2 from scratch (blue line).

At step 3, we investigate catastrophic forgetting [37, 42], by testing dataset 1 again. Especially in cases of increased transparency, catastrophic forgetting indeed occurs: the network substantially fails to remember the original training on MNIST+noise after being retrained on MNIST+cifar (Figure 2D) and vice versa (Suppl. Fig. 1D). We next show that increasing either the depth or the width of the network does not alleviate forgetting. For depth, we use the well-known VGG-16 [51]. For width, we increase twofold the number of filters or hidden units in the convolutional and linear layers, respectively. Figure 2E-G and Suppl. Figs. 1E-G, 2 show that the accuracy of the network in the *test CF condition* (orange line) is lower than in the *matched condition* (blue line). In summary, Figure 2B-G demonstrate that continual learning in both shallow and deep networks, and for either dataset order, gives rise to catastrophic forgetting.

### 3.3 Simple networks with contextual input, output, or *feedforward* switching units fail to perform sequential context switching

We next test three simple strategies that could avoid catastrophic forgetting. The first strategy is to add a binary contextual input to the NN in Figure 1E, similar to the input received by the recurrent neural networks in [36]. The input is added to the hidden layers (Suppl. Fig. 3A(a)) and is set to 0 for dataset 1 (e.g., MNIST+noise) and 1 for dataset 2 (e.g., MNIST+cifar). This strategy however is ineffectual: we find that catastrophic forgetting still occurs (orange vs blue lines in Suppl. Fig. 3C,F). Another strategy we tried was to require a separate binary output for context: 0 for dataset 1 and 1 for dataset 2. Again, this fails to reduce catastrophic forgetting (orange vs blue lines in Suppl. Fig. 3D,G). The idea underpinning these strategies was that by either directly providing knowledge about the identity of the context (noisy or CIFAR-10 context) – or explicitly requiring the network to represent this context in its output – the NN would be able to separate contextual information from digit information, and would learn to use only the latter in the classification task.

A third, related strategy takes inspiration from the biological V1 circuit discussed above [55, 65], where VIP neurons turn ON and OFF depending on an animal’s motion context. In our first, simplest model inspired by this circuit, we add *switching units* akin to the VIP neurons: they are OFF for the first context and ON for the second context. This architecture is the “ feedforward switching network”, with the NN from Figure 1E as the main network, while the switching units added are auxiliary units (Figure 1G, Suppl. Fig. 3A(b)); see the next section for a recurrent version of this architecture which give different results. The idea here is that after training on the first context with the switching units OFF, we can freeze learning on the weights from main network. In the *re-matched condition*, we train on the second context, while turning ON and only learning weights from the switching units, holding other parameters of the NN fixed. The contribution of the switching units gets added to the activations of main network hidden layer before it gets through a ReLU non-linearity. This is a similar network to the one with binary inputs described above, with one important difference: after training for dataset 1 with the switching units OFF, we are *freezing* the weights already learned and only changing the weights from the switching units.

For this feedforward switching network we are guaranteed to remember the first learned dataset by simply keeping the switching units inactive, while classification on the second dataset is enabled by turning ON the switching units. This strategy overcomes catastrophic forgetting if the network can solve the task on dataset 2 by only learning the weights from the switching units. As expected from computational work studying the complex recurrence of the V1 circuit [55], we find that this feedforward switching network achieves accuracies on dataset 2 (Figure 3C, green line) only slightly higher than the *unmatched condition* when using a network trained on dataset 1 alone (orange line), and far below the reference *matched* condition, when the network was trained on dataset 2 (blue line). As before, the conclusions hold when dataset 1 is either MNIST+noise or MNIST+cifar (Suppl. Fig. 3E), and we assume this is the case throughout the paper unless otherwise noted. In sum, while the feedforward switching network does not, by construction, show catastrophic forgetting, it has an inferior performance on the second context (dataset 2).

**Figure 3:**
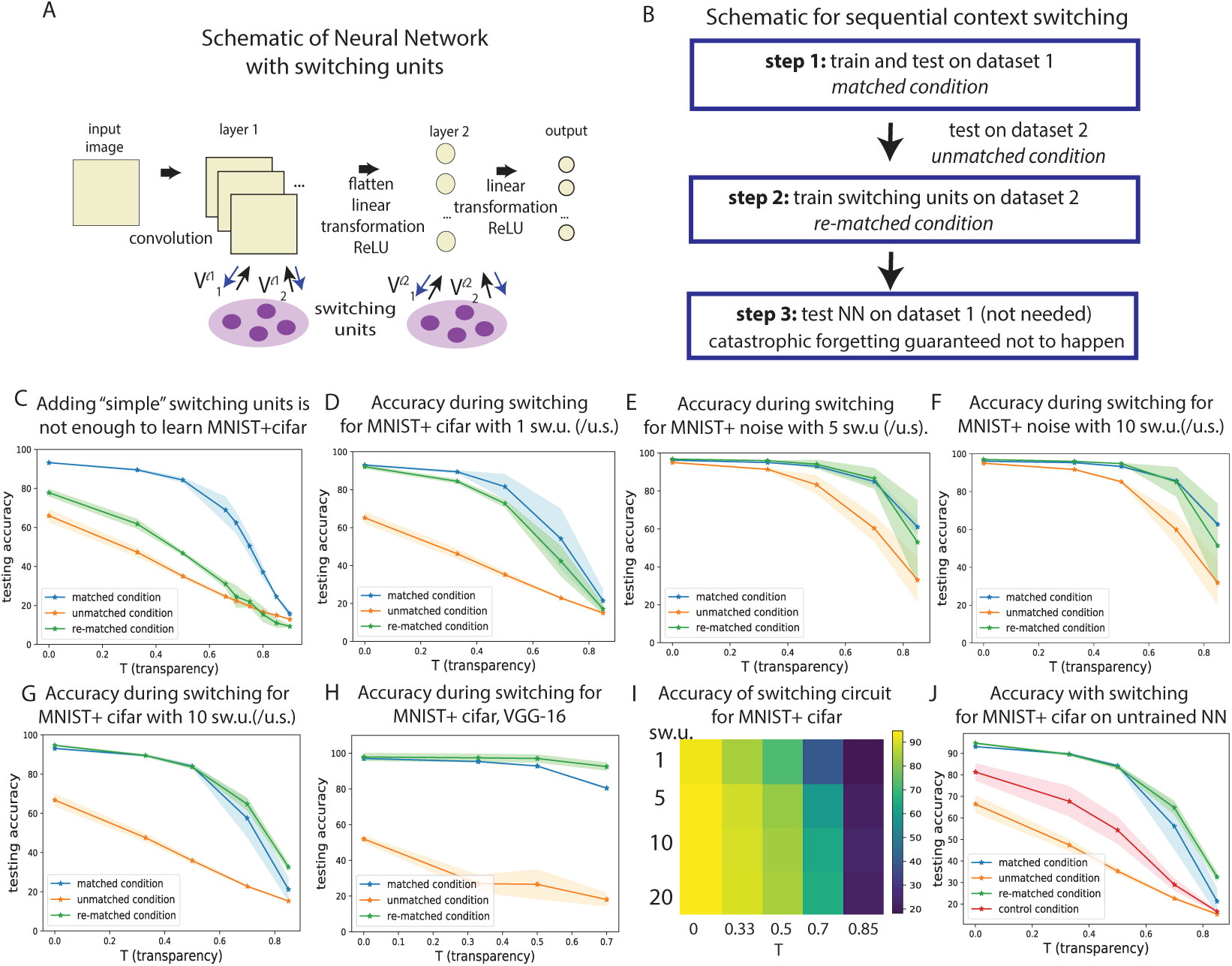
A). Schematic for the switching network. B). Schematic for sequential context switching: training and testing on datasets 1 and 2. C). Testing accuracy on MNIST+cifar vs transparency *T* : after training in the *matched condition* (step 1, blue line); after training in the *unmatched condition* (orange line); after training in the *re-matched condition* with switching units (per unit space for convolutional layers; step 2, green line). “ sw.u.” stands for switching unit(s), “ u.s.” stands for unit space. For D)-J) we always add the recurrent connections to the switching units and for D)-G) we test the corresponding dataset using 1, 5, or 10 switching units. H). Testing accuracy on the VGG-16 with switching units. I). Heat map of testing accuracy on the basic network across number of switching units and transparency *T*. J). Testing accuracy on the basic network (red line), when training weights to and from the switching units, but keeping weights in the main network randomly initialized. Blue, orange, and green lines are displayed for comparison.

We conclude that none of the three strategies considered above is successful in sequential context switching. In the next section, we introduce an improvement to the feedforward switching network that attains success.

### 3.3 Networks with *recurrent* switching units succeed in performing sequential context switching

We can improve the feedforward switching network by adding recurrent connections to the switching units (blue arrows, Figure 3A) [55]. When adding switching units to convolutional layers, we essentially add kernels, and we refer to switching units per unit space (sw.u/u.s.). The activities of this improved network – the switching network – at layer *l* for the second context *c*_2_ can be expressed as:

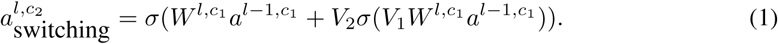

where 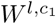 are the weights between layers *l*−1 and *l* of the main network for dataset 1 and first context *c*_1_, *σ* is the non-linearity (ReLU),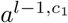 are the activations at layer *l*−1 for the first context *c*_1_, and *V*_1_, *V*_2_ are the weights to and from the switching units, as shown in Figure 3A.

When the switching units are ON, we learn connections to and from these units, with the goal of classifying dataset 2. This strategy succeeds: we find that the accuracy on dataset 2 for different values of *T* is much improved when adding even one switching unit (green line in Figure 3D). We can compare this performance to the *unmatched* and *matched conditions*: the *unmatched condition* entails testing dataset 2 on the NN trained on dataset 1 (orange lines) and represents a lower bound, while the *matched condition* implies testing dataset 2 on the NN trained on dataset 2 (blue lines) and is the accuracy we want to reach or exceed. As we add more switching units, the performance of the switching network on dataset 2 (*re-matched condition*) approaches or even surpasses the *matched condition*. Using 5 switching units, the performance is at times superior to the *matched condition* (Figure 3E), with an even more pronounced boost in performance as we increase the number of switching units to 10 (Figure 3G, p-value *<* 8.6· 10^*−*5^ on MNIST+cifar, and *T >* 0). When adding 10 switching units we use small 3 × 3 kernels, which causes the total number of parameters to be comparatively reduced. Switching units applied to the convolutional layers of the much deeper VGG-16 network also achieve sequential context switching, with superior accuracy in the *re-matched condition* using switching units to classify MNIST+cifar (Figure 3H, Suppl. Fig. 11).

We see that generally few switching units are required for the highest accuracy to be achieved. For the VGG-16, we use approximately a tenth of the number sw.u/u.s. present in the VGG main network. For the basic NN, the accuracy either plateaus or increases slowly, such that with 5 −10 switching units we are close to peak performance (Suppl. Fig. 4A-C). Figure 3I shows a summary of how testing accuracy in the basic network varies with the number of switching units and transparency, and we note the same tendency of the accuracy to plateau as switching units increase.

We then ran a series of controls to further test the effectiveness of the switching network. First, we altered dataset 2 to have shuffled labels, while dataset 1 had typically assigned labels (“ 1” for a written digit of 1, etc.). We find that switching units do not work as well when the context is unchanged, but the task has different input-output dependencies (Suppl. Fig. 4D,E). Second, we assessed a network where only weights to and from the switching units are learned, while the weights in the main network are random (Figure 3J, Suppl. Fig. 4F). The goal is to establish baseline performance when the main network does not contribute features from dataset 1 to the task. We find that dataset 2 accuracy barely surpasses the *unmatched condition*, especially when the switching network learns MNIST+cifar (red line, Figure 3J) and conclude that main network weights and activations from training on a similar dataset are necessary for sequential context switching.

We conclude that few switching units that are recurrently connected to the main network layers can provide substantial performance improvement in the sequential context switching task, and that main network activations, tuned to dataset 1, are necessary for this performance.

### 3.5 A comparison between the recurrent switching network and two established continual learning methods

We compare the switching network (with recurrent connections to switching units from now on) with two other learning methods that could achieve generalization across contexts: Elastic Weight Consolidation and Progressive Networks.

#### Elastic Weight Consolidation (EWC)

EWC [24] averts forgetting of old tasks by constraining learning on the weights important for those tasks. The importance of parameters for a particular task is quantified by the diagonal of the Fisher information matrix. The important weights stay close to their old values, keeping the parameters in a region of low error for task 1, centered around *W*_1_ (the weights for task 1) while learning task 2.

#### Progressive Network (ProgNet)

ProgNets [46] maintain a set of pre-trained networks (“ columns”) for each task and learn lateral connections between these columns to extract useful features from related tasks (Suppl. Fig. 6 for a schematic). During learning, the parameters of columns trained on previous tasks are kept constant, so weights are not overwritten and the network is immune to catastrophic forgetting by design. A drawback of this approach is the growth in the number of parameters with the number of tasks. We introduce another version of ProgNet, which we call “ ProgNet2” where the lateral connections from previous column layers are initialized to be the same as the feedforward connections between the corresponding column layers.

#### Performance comparison

We compare the performance of the switching network with the performance of four other networks, all having the same architecture as the main network (Figure 1E) of our switching network in common: the main network implementing EWC during learning (“ EWC network”); a ProgNet with two columns like the main network and lateral connections (“ ProgNet”); a network like ProgNet, save for how lateral connections are initialized (“ ProgNet2”); a main network with two separate last hidden layers for each context.

When networks are trained on the *matched condition* for MNIST+noise, they perform equally well since initially all of them use the same architecture (Figure 4A). The switching network and both ProgNets avoid catastrophic forgetting by design, but that is not the case for the EWC network. Varying *λ*, the trade-off hyperparameter that balances how well the network performs on dataset 2 versus how close the weights for dataset 1 and 2 are, we were unable to find a regime for EWC where both accuracies for the *test CF condition* and the *re-matched condition* were high. We conclude that, unlike the switching network, EWC cannot perform sequential context switching on these datasets at high transparency using this architecture. A possible explanation for this problem is the high complexity of the task compared to the complexity of the architecture used, which leads to a large fraction of the weights being essential for the first context, leaving few unimportant weights to learn the second context.

**Figure 4:**
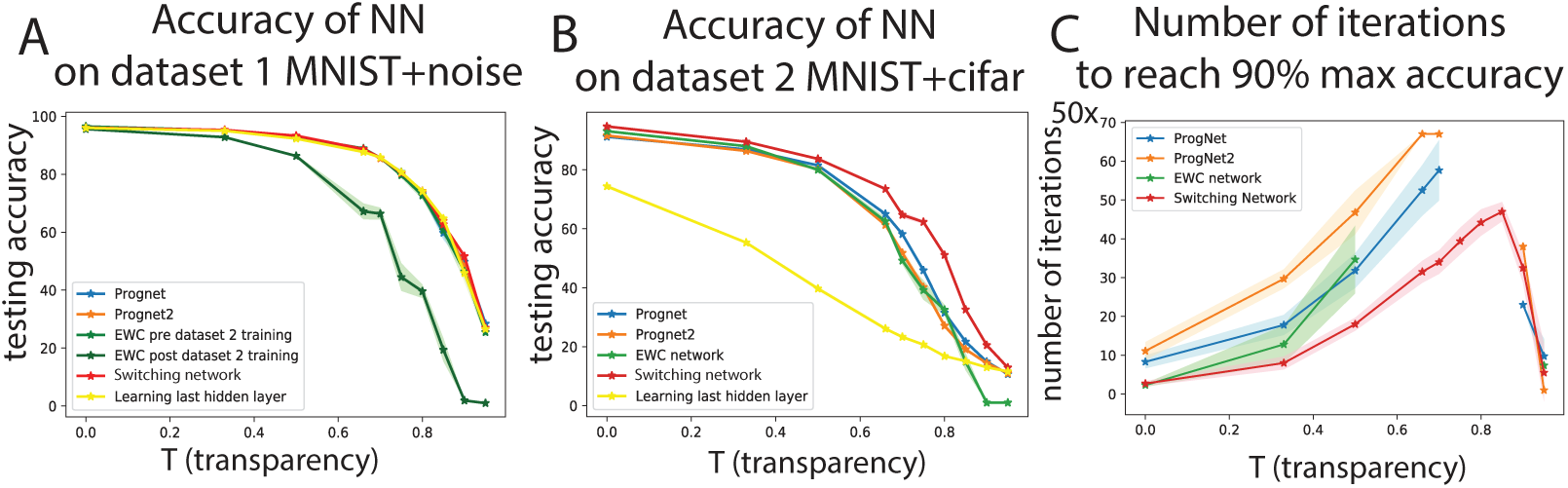
A) Testing accuracy for dataset 1 (MNIST+noise) for the five NNs. Only the EWC after training on dataset 2 (MNIST+cifar) shows a poorer performance because it cannot overcome catastrophic forgetting. B). Testing accuracies on dataset 2 (MNIST + cifar) for various NNs. We note that the switching network has a slight edge across all transparencies. C). Number of iterations (in steps of 50) for each NN, during training on MNIST+cifar, to reach 90% of the max. peak accuracy.

We next test on dataset 2 (MNIST+cifar), to find that the switching network performs slightly better than the ProgNets across all transparencies (Figure 4B). Furthermore, switching networks are faster than the ProgNets on transparencies ≤0.7 (Figure 4C), when considering the number of iterations to reach 90% of the peak accuracy for that transparency, maximized between the switching network, ProgNet, ProgNet2, and EWC. Several data points are not shown for ProgNet, ProgNet2, and EWC because the respective network does not reach 90% of this peak accuracy. The comparatively rapid increase to peak accuracy could be explained because the switching network has fewer weights to learn than the ProgNet, in addition to taking advantage of the features learned from dataset 1. Using 10 switching units for each layer (sw.u/u.s. with 3 × 3 kernels for the convolutional layers), we add an additional 10 × 3 × 3 × 2 + 10× 2 = 200 parameters to learn, compared to *O*(100, 000) for ProgNets. This over-parametrization might be the reason why for higher transparencies *T* these networks never reach the accuracy levels of the switching network, as the learning procedure could be stuck in a local minima. In conclusion, for most transparencies, our switching network shows enhanced performance at sequential context switching in terms of combined accuracy and learning speed (Figure 4, Suppl. Fig. 7).

### 3.6 Analysis of a switching network mechanism for context switching

Having established the switching network as a suitable architecture for sequential context switching, we ask what mechanisms underlie its performance. We first observe that switching units in the first layer have a predominantly negative effect on the activations of the main network, as seen from the histograms of contributions for switching to MNIST+noise and MNIST+cifar (Figure 5A). Importantly, the predominantly negative weight contributions from switching units were not built into the network, but emerged during learning on the second context. While this invites a direct comparison to the underlying biology, as VIP neurons are inhibitory, care and further study are required as VIP neurons can also have net disinhibitory effects through other pathways [65]. While the inhibitory dominance is not apparent at the second layer, we note that the switching units at the first convolutional layer are the main drivers of high performance (Suppl. Fig. 5).

**Figure 5:**
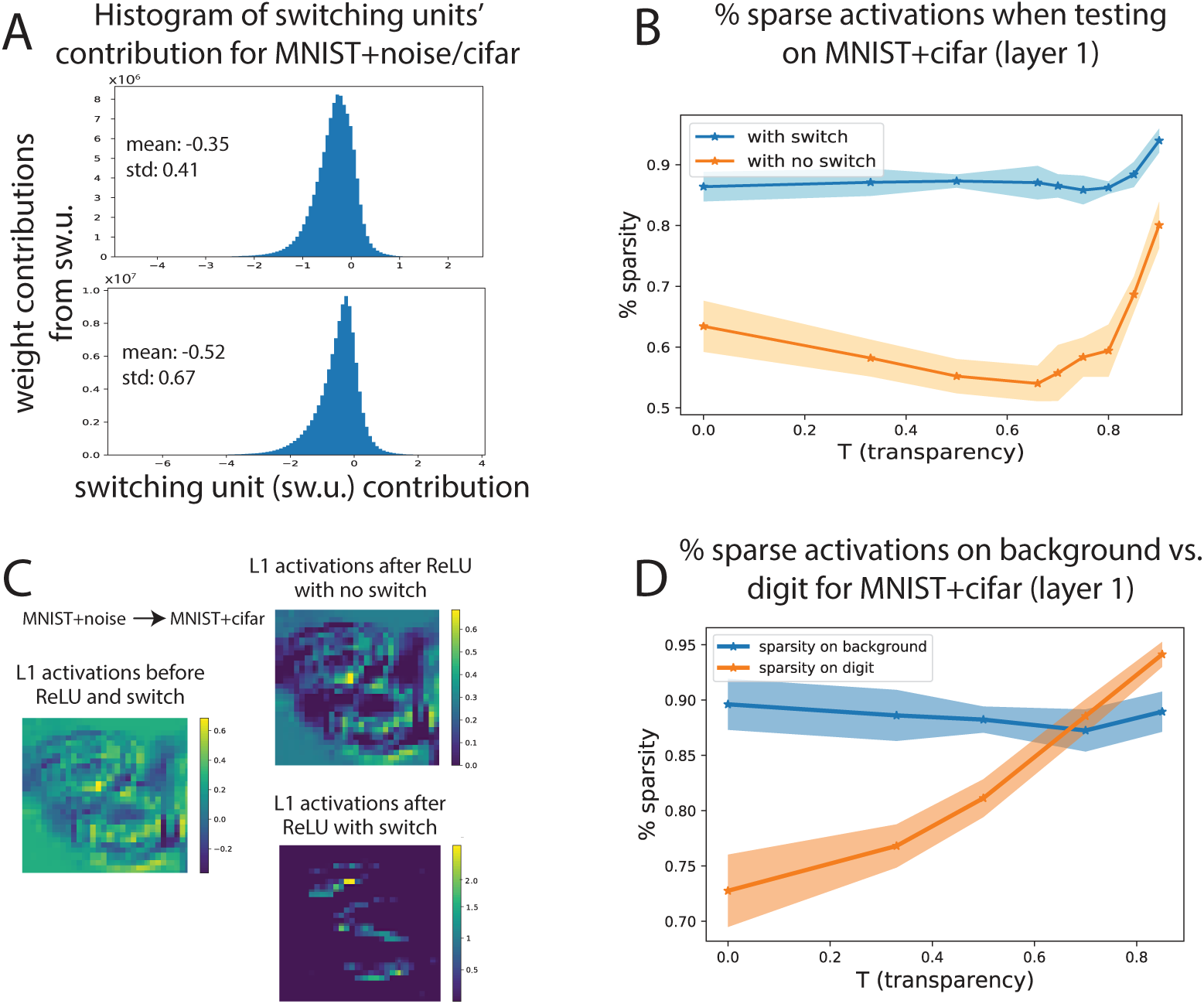
A). Histograms of switching unit contributions to the main network (for switching to MNIST+noise, up; for switching to MNIST+cifar, down). B). Percentage layer 1 sparse activations with or without adding the switching contributions (blue vs orange line), and after applying ReLU. C). Examples of (convolutional) layer 1 activation patterns with or without the switching contribution. D). Percentage layer 1 sparse activations within the background (blue) and within the digit (orange) when switching to MNIST+cifar after adding the switching contributions and the ReLU non-linearity.

We next make an allied observation about network representations. The negative contributions from switching units favor a sparsification of activations (after applying the ReLU non-linearity). Specifically, the activations after switching are much more sparse than those without switching (Figure 5B), and we hypothesize that, at least for lower transparencies where the background is the main nuisance, this allows the network to highlight features useful for digit classification while inhibiting redundant features from the background (Figure 5C, Suppl. Fig. 9). The validity of this hypothesis for lower transparencies is reinforced by evidence that the switching units inhibit a larger percentage of the background than of the digit (Figure 5D, Suppl. Fig. 9C), indicating that the background features are suppressed preferentially.

Finally, we observe that a low rank contribution can yield a similarly high performance (Suppl. Fig. 8) in a linearized version of the recurrent switching component of the model. This suggests that varied network mechanisms that implement low-rank updates to weight matrices may be sufficient to support contextual switches.

## 4 Conclusion

We study a bio-inspired switching network that is capable of domain adaptation between backgrounds with different statistical properties while the task is maintained. We construct a dataset which can be used as a benchmark for this task, by overlaying MNIST digits with different transparencies on noise or CIFAR backgrounds. The switching network is able to leverage features learned in one context and use them to do digit classification in a second context without forgetting classification on the first context.

While we do not have a precise mechanism on how the switching allows the network to perform well under both contexts with so few neurons, a set of analyses allow us to speculate. We did observe a sparsification of the representation with the switch on (Figure 5). We believe this allows the network to select the features relevant for the task; these features are different from the features in the new background, which are distractors. The fact that this can be obtained with a few neurons is backed by the analysis showing that a low rank change in connectivity matrix can result in a similar performance (Suppl. Fig. 8). A small number of neurons recurrently connected can provide the mechanism for a low rank matrix change in the connectivity needed for this task.

While we attain solid performance on a context-dependent image dataset, our findings have several limitations. First, the dataset is a simple benchmark with few categories (MNIST digits). We have opted for this dataset in order to remove other potential dataset confounds, but more complex datasets should be tested. Second, while we found high performance with relatively low numbers of switching units, especially for the VGG network, we do not currently have theoretical guarantees on the number of switching units sufficient for sequential context switching. Moreover, we have not studied transition between multiple contexts, although the suppression of the contextual features shown above opens the possibility that multiple context switches can be incorporated. Finally, the network does not include a downstream module to detect novel contexts. While the network can easily incorporate a module to detect contexts given *a priori* knowledge of the backgrounds (Suppl. Fig. 10), a future avenue of study will be to understand how such networks can detect new contexts in an unsupervised way.

In principle, switching networks employ a general circuit motif that implements a context-dependent computation, therefore switching units could be integrated to any neural network in order to address the problem of adaptation to context. A possible application area of high social importance is in addressing biases of background, in which networks can learn unwanted correlations between context and objects of interest [12]. Finally, this is one of few studies to address generalization across context in a bio-inspired architecture, while focusing specifically on background variation and removing other confounds (see [3]). In future years, we look forward to progress in understanding the context-dependent computations that enable the extraordinary versatility of biological agents to adapt and switch environments so effortlessly. Deciphering the general principles behind contextual generalization will hopefully enable our current networks to develop from being sophisticated “ pattern-matching machines” to more intelligent, human-like learners capable of abstracting visual concepts [3].

## 5 Acknowledgments

We gratefully acknowledge support of the Swartz Foundation Center for Theoretical Neuroscience at the University of Washington, and of NIH Training Grant 5 R90 DA 033461-08. We thank the Allen Institute for Brain Science founder, Paul G. Allen, for his vision, encouragement, and support.

