## Supplementary Material for "A biologically inspired architecture with switching units can learn to generalize across backgrounds"

### 1 Methods

#### 1.1 Dataset

To address the problem of image classification with context switching, we created a dataset whose only confounding feature is context. We make this dataset publicly available under the Creative Commons v 4.0 license as a benchmark to test generalization across context: https://figshare.com/s/e5807249c28ec5bcff94.

The dataset consists of $32 \times 32$ sized images of MNIST digits set on either noisy backgrounds, with background pixel intensity drawn independently from Gaussian random variables, or MNIST digits set on a more naturalistic background from the CIFAR-10 dataset. We refer to the subset of MNIST digits set on noisy backgrounds as the "MNIST+noise" dataset and to the subset of MNIST digits set on CIFAR-10 backgrounds as "MNIST+ cifar" dataset.

To construct these two sets of images, we first consider the normalized $28 \times 28$ pixel images from the MNIST dataset with maximum pixel value of 1, and refer to this normalized set unless otherwise specified. We then identify the pixels in these images that belong to the digit. The background in the MNIST dataset images is uniformly 0 therefore we consider all non-zero pixels to belong to the digit as opposed to the background. The pixels that constitute the digits are retained in a mask image to be used on the background, as described below.

For MNIST+noise, we create $32 \times 32$ pixel images where the pixels are the absolute value of Gaussian random variables of mean 0 and standard deviation 1, then normalize appropriately so that the maximum value is 1. The mask described above is super-imposed on this noisy background, after being properly shifted so that the digit pixels represented by the mask are in the middle of the noisy image (given that MNIST images are $28 \times 28$ and noisy image backgrounds are $32 \times 32$, there will be a 2 pixel shift). While pixels outside the mask are not modified, pixels from the noisy background that correspond to the mask (and hence digit) are linearly combined with MNIST digit pixels.

$$p_i = T \cdot p_i^b + (1 - T) \cdot p_i^{digit} \tag{1}$$

where $p_i$ is pixel $i$ from the MNIST+noise image, $p_i^b$ is pixel $i$ from the noisy background and $p_i^{mnist}$ is pixel $i$ from the MNIST image. $T$ is a parameter that makes the task more difficult, and we refer to $T$ as digit *transparency*. As we increase the transparency, the background interferes with the digit more and the identity of the digit becomes more ambiguous.

For MNIST+cifar, we consider the $3 \times 32 \times 32$ images from CIFAR-10 and average across the color dimension, then normalize. We then proceed analogously as for MNIST+noise to create MNIST+cifar, a dataset of MNIST digits on more naturalistic CIFAR-10 backgrounds. Each MNIST digit is super-imposed on a unique CIFAR-10 background, so that there is a one-to-one correspondence between the image digits and backgrounds (although this is not important). The correspondence is random, ignoring the CIFAR-10 class corresponding to the background image. The number of samples in the CIFAR-10 dataset, for both training and testing, is smaller than in the MNIST dataset, therefore we limit the number of samples in our MNIST+cifar dataset to that of the CIFAR-10. We limit the number of samples in the MNIST+noise data in a similar way so that the number of samples in the two datasets is identical.

The goal of these two datasets is to perform image classification and correctly identify the MNIST digit despite the different interfering backgrounds.

### 1.2 Experiments

We carried out all experiments using the Torch neural network framework on two machines as described below in the *Computing resources* subsection. We used the dataset with $32 \times 32$ MNIST digit images set on different backgrounds described above, except for the experiments on the deeper network VGG-16 [5], when we re-sized these images to be $224 \times 224 \times 3$ by first using the resize method with inter-area interpolation from the cv2 python library, and then concatenating two other $224 \times 224$ identically 0 matrices.
We split both the MNIST+noise and the MNIST+cifar data into 80% training and 20% testing samples. When using a validation dataset (either for determining the number of switching units to be chosen or to search for the hyperparameter $\lambda$ in the case of the EWC algorithm), the split is 64% training, 16% validation, and 20% testing data from the total samples. This partition of training and testing data does not correspond to the classic MNIST data training and testing split. Throughout all experiments we considered both the case when the order of training was MNIST+noise, then MNIST+cifar and vice versa.

#### 1.2.1 Network architectures

We tested both simple (toy) neural network (NN) examples and a more complex, deeper architecture, the VGG-16 [5]. The *basic* network architecture used throughout the paper is shown in Figure 1E. Aside from the input and output layers, the network has 2 hidden layers, where the first is a $16 \times 28 \times 28$ convolutional layer and the second layer is a 64 unit fully connected hidden layer. The first convolution has a kernel size of 5, stride 1, 16 filters, no padding, and is followed by a ReLU non-linearity. The second operation is linear, between the flattened 12544 unit convolutional layer and the 64 unit hidden layer. After another ReLU non-linearity, the final operation is a $64 \times 10$ linear operation. All operations have biases in addition to weights.

There are two more networks we use to test the $32 \times 32$ MNIST+noise and MNIST+cifar datasets (Suppl. Figure 5). The second network we employ is a 3 hidden layer network with two convolutional layers and a fully connected layer. The network has the same architecture as before (kernel size 5, stride 1, 16 filters for the first convolution; 64 units for the fully connected layer), except for the second convolution which now employs 8 filters, with the same kernel size and stride. As before, these layers apply a ReLU non-linearity.
Another architecture is a 4-layer network with 2 convolutional hidden layers and 2 fully connected hidden layers. This network has a $16 \times 28 \times 28$ convolutional first layer, a $32 \times 14 \times 14$ convolutional second layer (using a $5 \times 5$ kernel and stride 1 as before), an 128 unit fully connected layer, and a 64 unit fully connected layer. In addition to the ReLU non-linearity, the network uses max-pooling after every convolution and dropout with $p = 0.2$ for all except the last hidden layer.
These two additional networks are tested to assess the effectiveness of switching units on various architectures, and at different layers. We find that switching units are most useful when added at the first convolutional layer, although there is some benefit, albeit smaller, to adding switching units at later hidden layers (Suppl. Figure 5). These findings lead us to apply switching units only at the first convolutional layers of VGG-16.

In the case of VGG-16, we replaced the final layer with two fully connected hidden layers with 256 and 10 units respectively, the later corresponding to the number of classes in our dataset. Regarding switching units, we added about a tenth of the number of units to the corresponding hidden layer: 6 switching units per unit space (sw.u./u.s.) for the 64 filter hidden layer; 12 sw.u./u.s. for the 128 filter hidden layer; 50 sw.u./u.s. for the 512 filter hidden layer. We only added switching units to six out of the thirteen convolutional layers, and specifically to the first three layers. No switching units were added to change activations of the latter linear layers.

#### 1.2.2 Training the networks

For the basic network (Figure 1E) and for the other 3− and 4− hidden layer networks, we used the ADAM optimizer, with variable learning rate that was not a priori fixed, and cross-entropy loss (Table 1 below). For the simple network enforcing a contextual output (Suppl. Fig. 3A(a)) we used a cross-entropy loss for the output layer units doing digit classification and a mean squared error (MSE) loss for the contextual output. For the simple one layer network for context classification where the binary output indicates whether the background is noisy of CIFAR-10 (Suppl. Figure 10), we use the MSE loss.

For VGG-16, we use the network pre-trained on ImageNet. During training of the main network, we keep the first eleven (out of thirteen) convolutional weights fixed while varying the other weights in the network. We tried a variety of optimizers with different hyperparameters, with distinct results depending on whether dataset 1 was MNIST+noise or MNIST+cifar. After a thorough search, see Table 1 below for a summary of all hyperparameters used during training.

| **hyperparameters** | basic network/ switching network | VGG-16, MNIST+noise | VGG-16, MNIST+cifar |
|---|---|---|---|
| optimizer | ADAM | RMSProp | SGD |
| learning rate | – | 0.0001 | 0.01 |
| other hyperparameters | – | – | momentum= 0.9, weight decay= $5 \cdot 10^{-4}$ |
| loss function | cross entropy | cross entropy | cross entropy |
| epochs | 5 | 5 | 5 |
| batch size | 50 | 32 | 32 |

**Table 1:** Hyperparameters and optimizers used for training MNIST+noise and MNIST+cifar on the basic network, switching network, and VGG-16 with switching units.

These hyperparameters were chosen to generate Figures 2 and 3, and the Supplementary Figures.

We mention that for VGG-16 the weights to and from the switching units were learned with RMSProp using a learning rate of 0.0001, momentum 0.9, and weight decay $5 \cdot 10^{-4}$, whether learning MNIST+noise or MNIST+cifar. These were chosen after careful experiments testing different learning methods and hyperparameters.

We attempt to use a validation dataset in order to choose an optimal number of switching units. Predictably, as the number of switching units increases, the testing accuracy increases as well, however the increase is negligible and effectively the accuracy plateaus as seen in Figure 4A-C. For a majority of the experiments, when not specifically indicated, we use 10 switching units (per unit space with a kernel size of 3 for convolutional layers) for the switching network, as this adds a limited number of parameters while maintaining high accuracy. We add a fixed number of switching units to the VGG-16 as described above in the *Network architectures* section (about a tenth of the corresponding hidden layer units).

The switching network may be equipped with a "context detector". We trained a simple one layer neural network to classify the backgrounds of images, whether it was noisy of CIFAR-10 backgrounds. The output of this network then determined whether the switch was ON or OFF during testing of a dataset with randomly interleaved MNIST+noise and MNIST+cifar digits (Suppl. Figure 10). We find that having the switching units turn ON or OFF depending on context (*adjusted switch*) gives higher performance than having the switch always OFF (*never switch* or *unmatched condition*) or having the switch always ON (*always switch*). The design of a network that detects new contexts in an unsupervised way is left for future work.

The results of experiments performed on VGG-16 using switching units are shown in Suppl. Figure 11A,B. It is worth highlighting the superior performance of the *re-matched condition* as compared to the *matched* and *unmatched conditions*: the switching network leverages the features learned by VGG-16 on dataset 1 to increase performance on dataset 2, with performance surpassing that of a network explicitly trained for dataset 2.

We run each experiment 10 times with different seeds for the basic network (and also for the 3- and 4- hidden layer networks), but only 5 times for VGG-16 because of the prohibitively long training time for this deep network. Error bars in the plots quantify the uncertainty in the test set and represent the standard deviation of the samples obtained from these experiments. We mention that uncertainty in Figure 2 and Suppl. Figure 1 is only w.r.t. network initialization, as there is no shuffling of the training data for each epoch/experiment; this can cause error bars to be comparatively small.

#### 1.2.3 Training using other continual learning methods

We compared our switching network with two prominent continual learning methods: elastic weight consolidation (EWC, [3]) and progressive networks (ProgNets, [4]). To implement these methods, we use two Github repositories: [1] for EWC, and [2] for ProgNet. Both repositories are under the MIT license, so we were able to appropriately change the code to suit our sequential context switching problem. Scripts in [2] underwent only minor modifications, whereas scripts in [1] had to be more substantially modified to be operational.

**EWC.** EWC is a continual learning method that protects consolidated knowledge through regularization. It takes inspiration from synaptic consolidation in the neocortex, where a proportion of synapses becomes less plastic and hence more stable over longer timescales in order to durably encode information. In EWC, each weight is pulled back toward its old values by an amount proportional to its importance in previously learned tasks, which slows down learning on these important weights. Effectively, EWC constrains important parameters to stay close to their old values and this constraint is implemented as a quadratic penalty (Equation (2)).
The posterior $p(\theta|D_A)$ is the probability of parameters $\theta$ given task data A ($D_A$) and contains information about which parameters are important for the task. We can approximate this posterior as a Gaussian distribution with mean given by the old values of $\theta$ on task A, $\theta^*_A$, and a diagonal precision given by the diagonal of the Fisher information matrix $F$.

Given that we consider the diagonal of the Fisher information matrix to be a good estimate of how important each parameter is, we can express the loss function at the second task B (after first task A) as:

$$L(\theta) = L_B(\theta) + \sum_i \frac{\lambda}{2} F_i(\theta_i - \theta^*_{A,i})^2 \tag{2}$$

where $L$ is the loss function that prevents catastrophic forgetting (CF), $L_B$ is a loss function on task B without considering CF, $\lambda$ sets how important the old task is compared with the new one, $i$ labels each parameter, $F_i$ is the $i$-th entry of the Fisher information matrix diagonal corresponding to parameter $\theta_i$, and $\theta^*_{A,i}$ is the solution found for the $i$-th parameter when learning task A.

We applied EWC on the main network (Figure 1E) used for the switching network. We used cross entropy error, the ADAM optimizer with variable learning rate, and fixed hyperparameters like the fisher estimation sample size (the number of samples used to compute the diagonal of the Fisher information matrix) to 1024. Using the validation set, we do a systematic search over $\lambda$, from values of 100 up to $10^{10}$ in steps of $10^i/2$, $i = 2, ..., 10$. We find that for high values of $\lambda$ ($\approx 10^8$), CF is indeed averted, but at the expense of lower accuracy for dataset 2. For lower values of $\lambda$, the accuracy on dataset 2 improves and is competitive with the switching network accuracy, but at the expense of forgetting dataset 1. Allowing different values of $\lambda$ for different transparencies did not overcome this issue. We conclude that, barring attempts to change the optimization hyperparameters, loss function, or fisher estimation sample size, EWC is unsuccessful at sequential context switching.

**ProgNet** The main idea of ProgNets is that CF is prevented by instantiating a new neural network – "a column" – for each task being solved, while transfer is enabled via lateral connections from features of previously learned columns. The parameters for all previous columns (and tasks) are held fixed while training the new column so that there is no interference between tasks and hence the ProgNets are immune to CF by design. A schematic of the ProgNet is shown in Suppl. Figure 6.

A ProgNet starts with a single column: in our case, the main network of Figure 1E used for EWC and for the switching network. We can denote the hidden activations of this column as $h_i^{(1)} \in \mathbb{R}^{n_i}$, where $n_i$ is the number of units at layer $i \leq L$ and parameters $\theta^{(1)}$ are trained for convergence. When learning on the second task, the parameters $\theta^{(1)}$ are "frozen" and a new column with parameters $\theta^{(2)}$ is instantiated with random initializations. Layer $h_i^{(2)}$ receives input from both $h_{i-1}^{(2)}$ and $h_{i-1}^{(1)}$ via the lateral connections. We can generalize to $K$ tasks so that the activations at column $k$ can be expressed as:

$$h_i^{(k)} = f(W_i^{(k)} h_{i-1}^{(k)} + \sum_{j<k} U_i^{(k:j)} h_{i-1}^{(j)}), \tag{3}$$

where $W_i^{(k)} \in \mathbb{R}^{n_i \times n_{i-1}}$ is the weight matrix of layer $i$ and column $k$, $U_i^{(k:j)} \in \mathbb{R}^{n_{i-1} \times n_i}$ are the lateral connections from layer $i-1$ of column $j$ to layer $i$ of column $k$, $h_0$ is the network input, and $f$ is an element-wise non-linearity.

An advantage of this approach is that ProgNets make no assumption about the relationship between tasks. Hence, columns in ProgNets are free to re-use, modify, or ignore previously learned features from columns via lateral connections. A downside of this approach however is the growth in the number of parameters with the number of tasks. Even for our two columns corresponding to MNIST+noise and MNIST+cifar, there are $O(100,000)$ additional parameters added for the lateral connections. As suggested in [4], only a fraction of the new capacity is probably utilized, which indicates that pruning or online compression during learning could alleviate this overparametrization.

For our training we use the ADAM optimizer with variable learning rate and the cross entropy loss. Our ProgNet has two columns, one trained on the MNIST+noise dataset, the other trained on the MNIST+cifar dataset. We probe sequential context switching in either order, as we did for EWC and the switching network. A version of our network which we call "ProgNet2" initializes before training the lateral connections between $h_{i-1}^{(1)}$ and $h_i^{(2)}$ with the feedforward connections between $h_{i-1}^{(1)}$ and $h_i^{(1)}$. Otherwise, these weights are randomly initialized for the standard ProgNet.

### 1.3 Computing resources

For our experiments we used two machines. One had two NVIDIA TITAN X (Pascal) 12G GPUs and a 20-core Intel(R) Xeon(R) CPU E5-2698 v4 with 512 GB RAM. The operating system used was Linux Ubuntu 18.04 LTS. The second machine had four GF RTX 2080 Ti GPUs and a 40-core Intel Gold 6230 CPU with 512 GB RAM.

Our (toy) switching network using the basic NN takes about 4 hours to train for a particular combination of switching units (e.g. 10 sw.u.(/u.s.)), for $T \in \{0.0, 0.33, 0.5, 0.66, 0.7, 0.75, 0.8, 0.85, 0.9\}$, and for 10 different seeds. Plotting Suppl. Figure 4C takes about 3 days. For VGG-16 it takes about 10 hours to train (for one seed), for a particular number of switching units chosen and for $T \in \{0.0, 0.33, 0.5, 0.7, 0.85\}$.

### 1.4 Code

A repository with the code to generate the figures in the main paper can be found at https://github.com/dvoina13/switching_network_ANN

### 2 Supplementary Figures

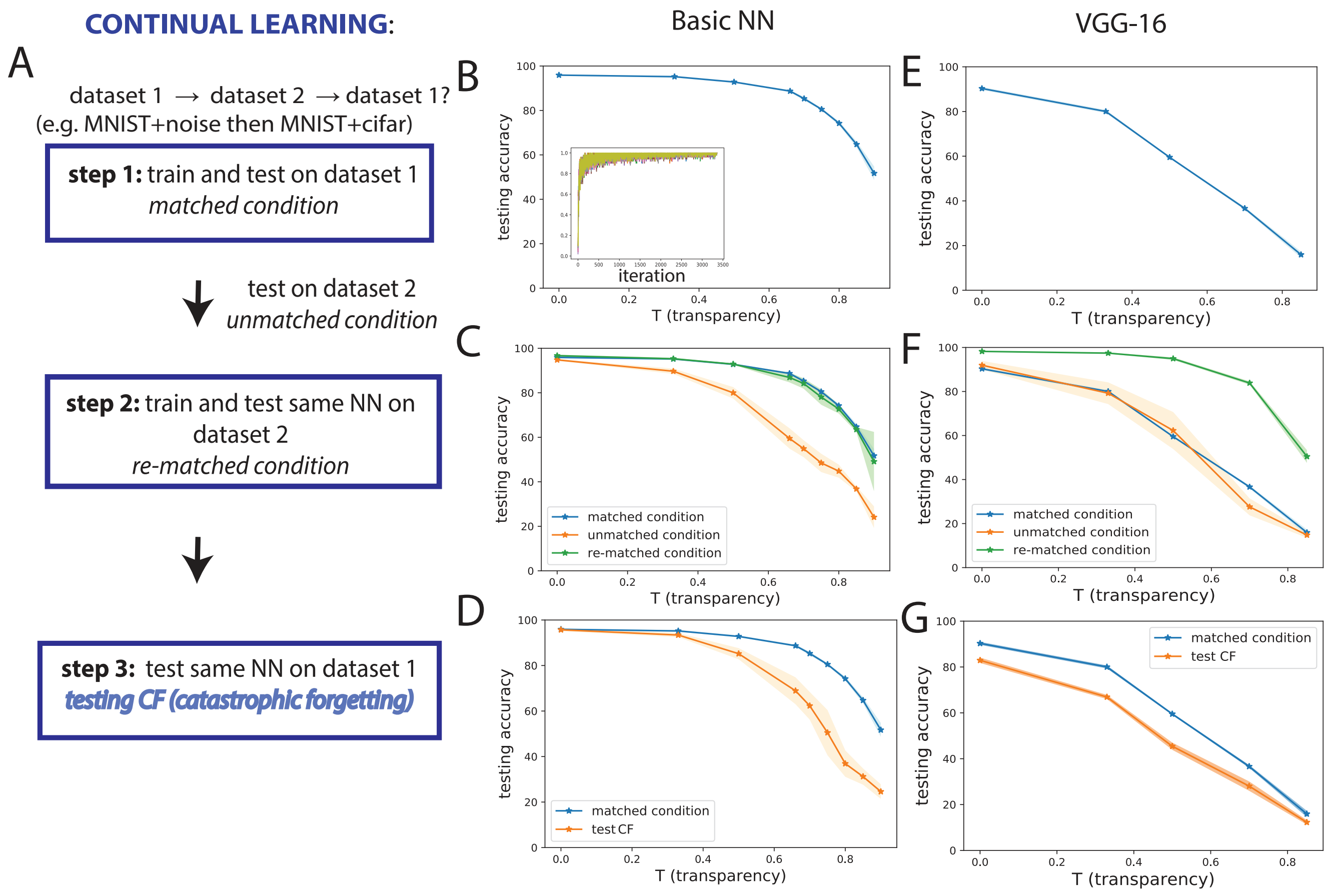

**Figure 1:** A). Schematic of training and testing for sequential context switching to test catastrophic forgetting. We first train and test on dataset 1 (Step 1), then train and test the same network on dataset 2 (Step 2), then finally test again on dataset 1 to probe forgetting (Step 3). B). Average accuracy for the *matched condition* on the MNIST+noise dataset as transparency $T$ increases (inset: convergence during training on the MNIST+noise dataset). C). The MNIST+noise dataset is less accurately classified by a network trained on MNIST+cifar (*unmatched condition*, orange line) than one trained on MNIST+noise (*matched condition*, blue line). However, once we re-train the MNIST+cifar-trained NN on MNIST+noise (Step 2), the accuracy approaches that for the *matched condition* (*re-matched condition*, green line). D). The accuracy when re-testing MNIST+noise on a network that was first trained on MNIST+noise, then on MNIST+cifar is reduced (*test CF condition*, orange line) compared to the *matched condition*, therefore we conclude catastrophic forgetting occurs. E)-G) same as B)-D) using the VGG-16 network instead of the basic network. This differs from main text Figure 2 in that catastrophic forgetting on MNIST+noise is tested (vs. MNIST+cifar).

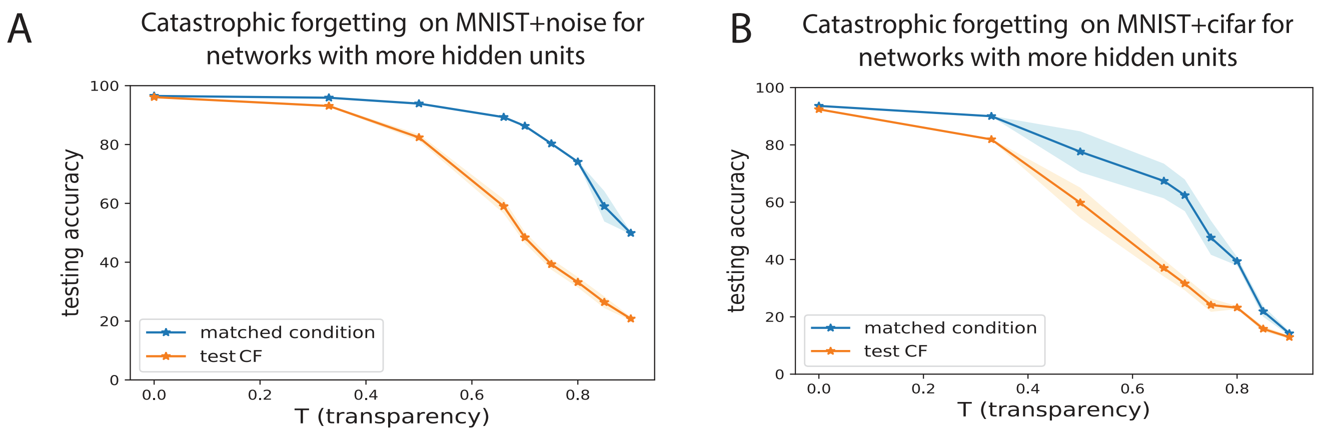

**Figure 2:** Increasing the depth of the network does not alleviate forgetting. To probe CF on wider networks, we increased twofold the number of filters or hidden units in the convolutional and linear layers, respectively. A). Accuracy on the MNIST+noise dataset for the *test CF condition*, after training on MNIST+noise, then on MNIST+cifar is lower particularly for higher $T$ than for the *matched condition*. This shows CF occurs. B). Same as in A), for the MNIST+cifar dataset.

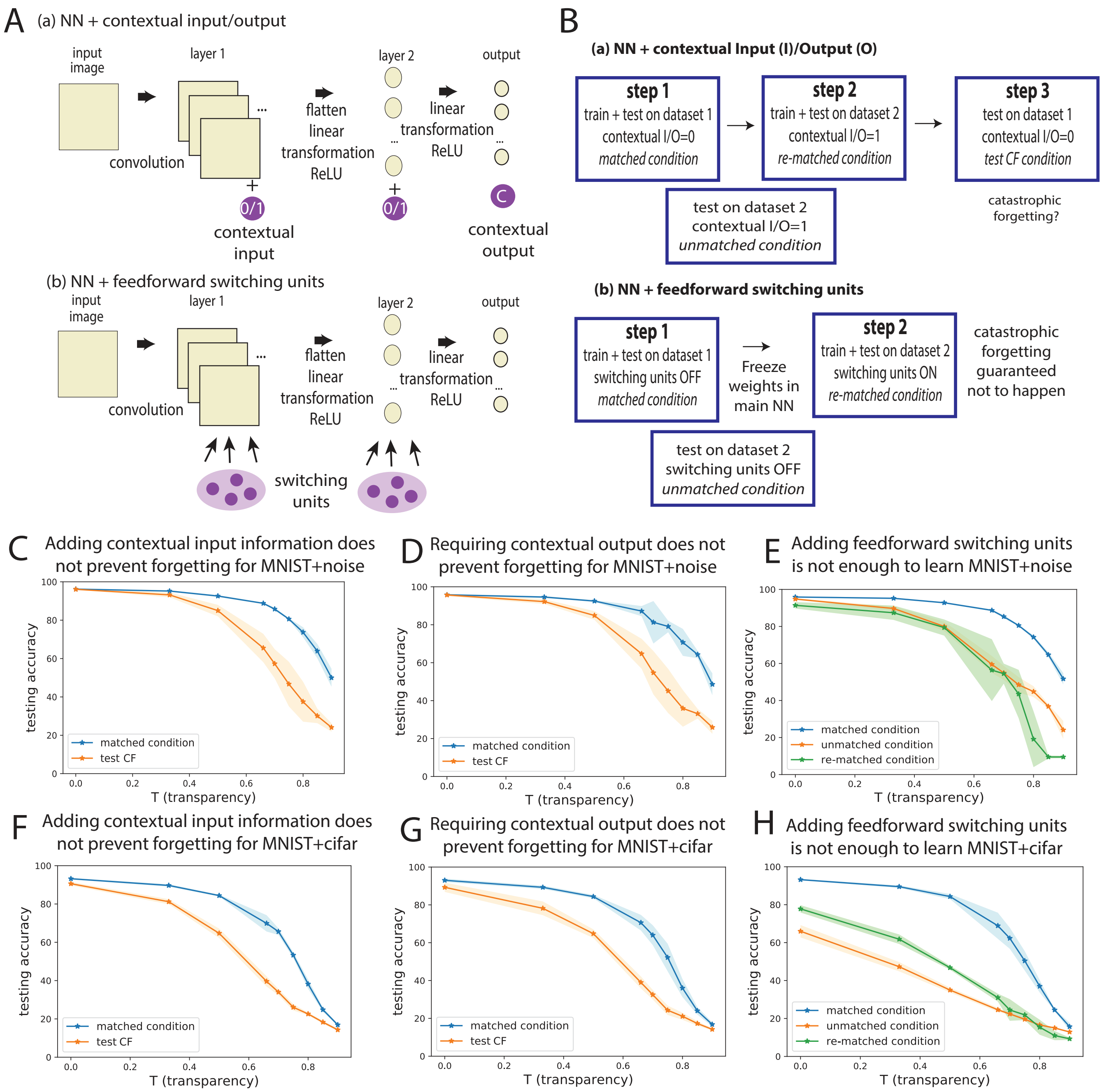

**Figure 3:** A). We test NNs that apply distinct strategies in order to overcome catastrophic forgetting. In (a), we either add a binary input to each layer with information about context (0 for dataset 1 and 1 for datatset 2), or we train the NN to output a binary variable signifying context. In (b), we add feedforward switching units to our NN. These units are OFF for dataset 1, and ON for dataset 2. Note that these units are different in structure compared to those described in Section 3.4 as they lack recurrent feedback from the corresponding layer they act upon. B). Schematic of how sequential learning is implemented for each NN described in A). C). The NN with contextual input does not overcome catastrophic forgetting. The accuracy on MNIST+noise in the *matched condition* (step 1) is significantly higher for certain T than the accuracy on MNIST+noise in the *test CF condition* (step 3). D). The NN with contextual output does not overcome catastrophic forgetting. Requiring a binary output still has the *matched condition* accuracy on MNIST+noise (step 1) significantly higher for certain T than the *test CF condition* accuracy (step 3). E). The NN with feedforward switching units added does not learn the classification task for dataset 2 at high accuracy. The accuracy in the *re-matched condition* for MNIST+noise, when training weights from the switching units, is not considerably improved from the *unmatched condition*, and considerably poorer than for the *matched condition.* F)-H). Same as in C)-E), but for MNIST+cifar.

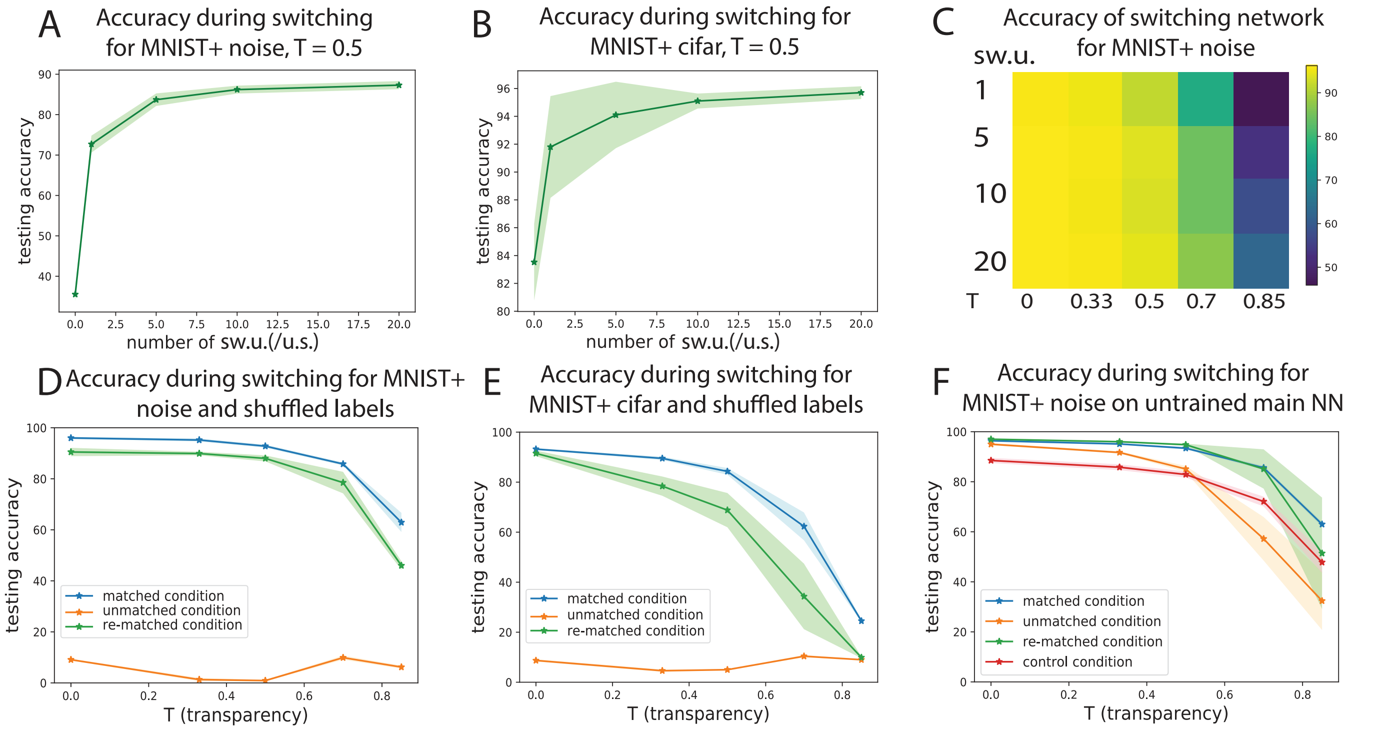

**Figure 4:** A). Accuracy for the switching network on MNIST+noise with $T = 0.5$, while varying the number of switching units (per unit space –sw.u/u.s.– for convolutional layers). B). Accuracy for the switching network on MNIST+cifar with $T = 0.5$, while varying the number of sw.u.(/u.s.). C). Heat map of accuracy on MNIST+noise for the switching network across number of switching units and transparency $T$. D). Accuracy for the switching network on MNIST+noise as labels are shuffled and the network must adapt to a new context as well as different input-output dependencies. This is a control experiment to show the switching network becomes less effective as tasks/datasets become more distinct. E). Same as D), for MNIST+cifar. F). Accuracy for the switching network on MNIST+noise (red line), when training weights to and from the switching units, but keeping weights in the main network randomly initialized.

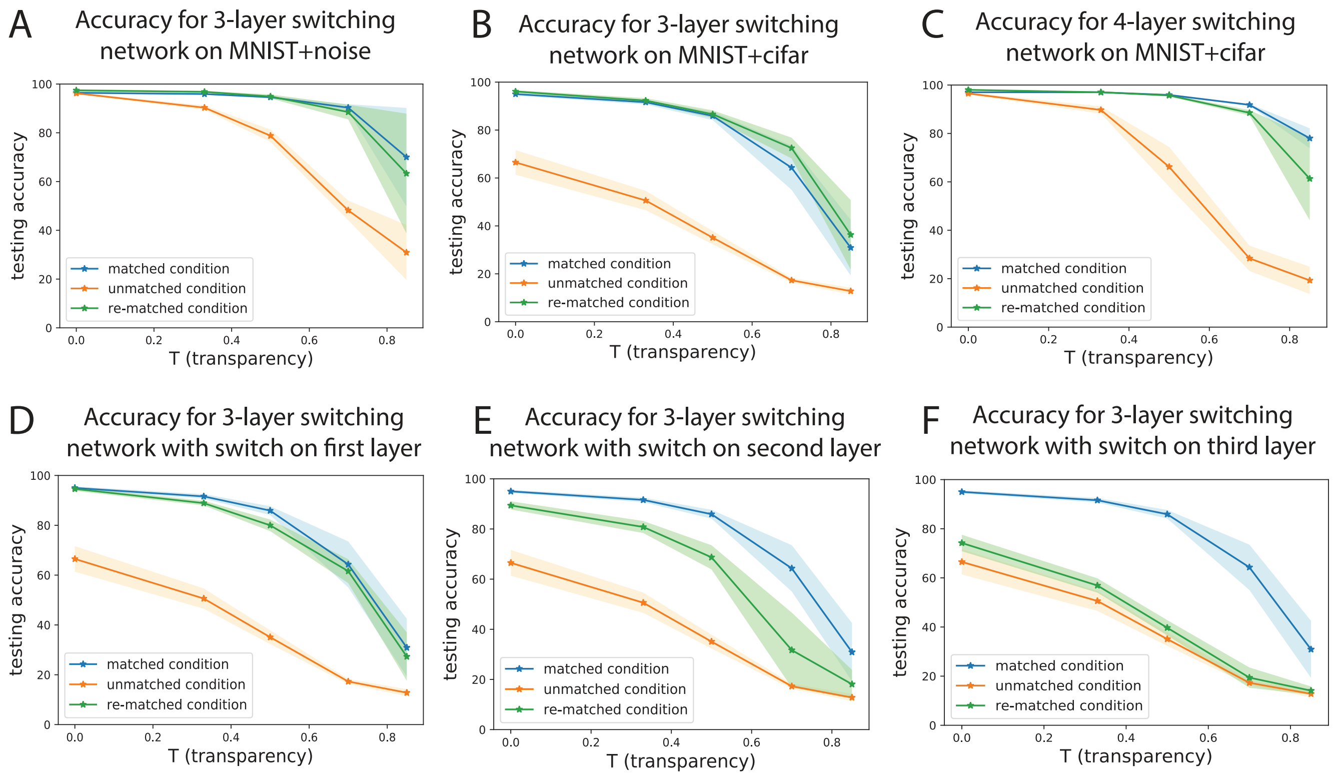

**Figure 5:** Switching networks with 3 and 4 hidden layers are effective. A). Accuracies on MNIST+noise for a 3 hidden layer NN as described in Section 1.2.1, *Network architectures*. B). Same as A), on MNIST+cifar. C). Accuracies on MNIST+noise for a 4 hidden layer NN as described in Section 1.2.1, *Network architectures*. D). Accuracies on MNIST+cifar for a 3 hidden layer NN when switching units are added only to the first hidden layer. We note that the switching architecture is most effective in this case. E). Accuracies on MNIST+cifar for a 3 hidden layer NN when switching units are added only to the second hidden layer. F). Same as D-E), but with switching units added only to the third hidden layer.

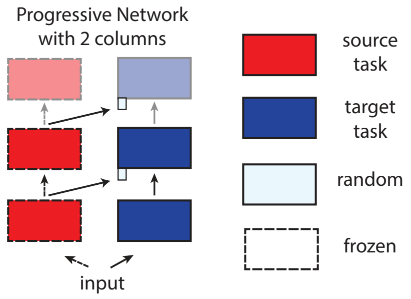

**Figure 6:** Schematic of ProgNets with two columns for two tasks. Reproduced from [4].

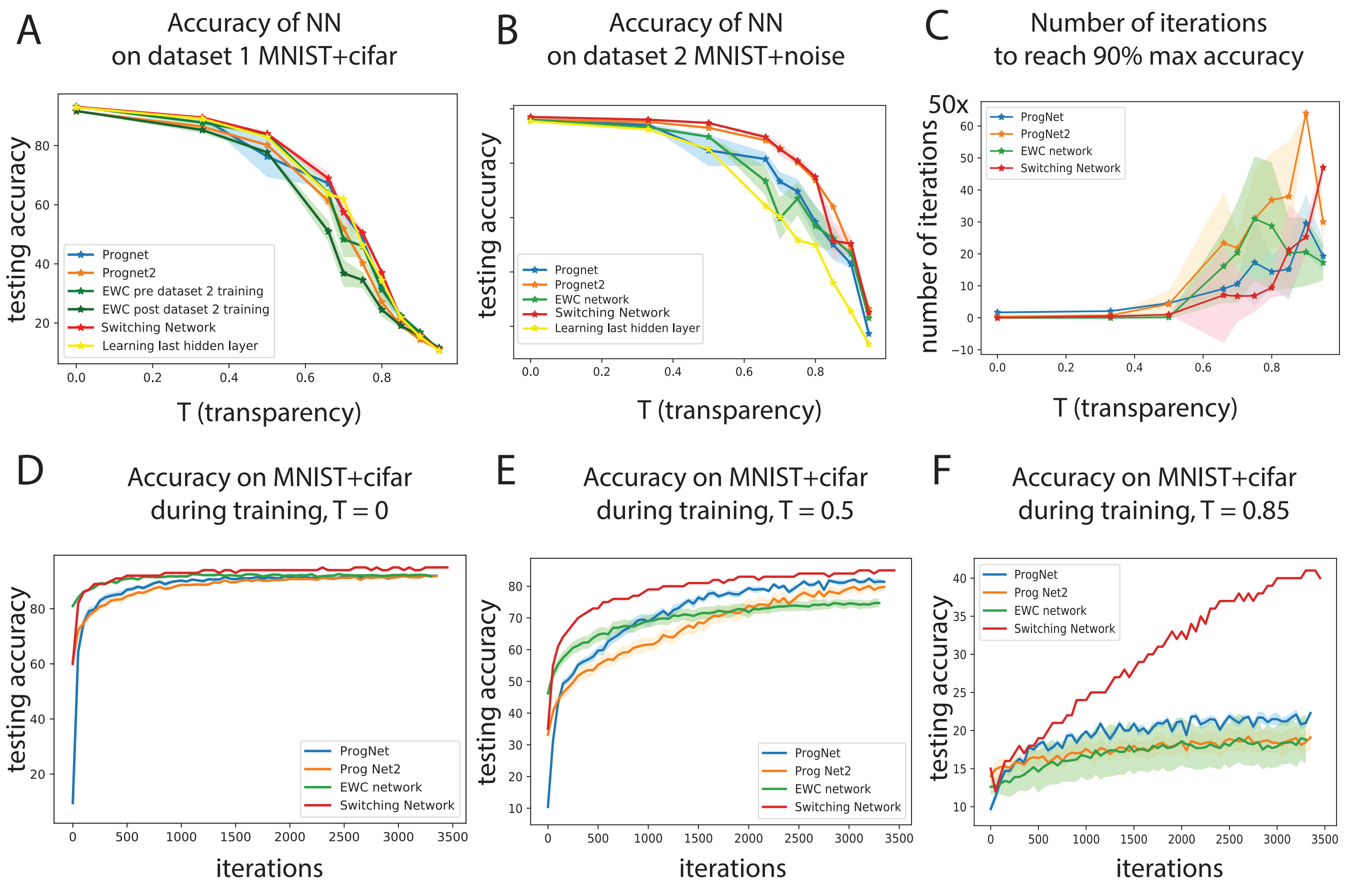

**Figure 7:** A). Accuracy on dataset 1 (MNIST+cifar) for the five NNs. B). Accuracies on dataset 2 (MNIST + noise) for various NNs. We note that the switching network has a slight edge across all transparencies. EWC achieves a poor balance between dataset 1 accuracy in the *test CF condition* (Suppl. Fig. 7A) and dataset 2 accuracy in the *re-matched condition* (Suppl. Fig. 7B). C). Number of iterations (in steps of 50) for each NN, during training on MNIST+noise, to reach 90% of the max. peak accuracy. D). Testing accuracy during training on MNIST+cifar with $T = 0$. E). Same as D), on MNIST+cifar with $T = 0.5$. F). Same as D), on MNIST+cifar with $T = 0.85$. A)-C) differs from Figure 4 in that here dataset 1 is MNIST+cifar and dataset 2 is MNIST+noise.

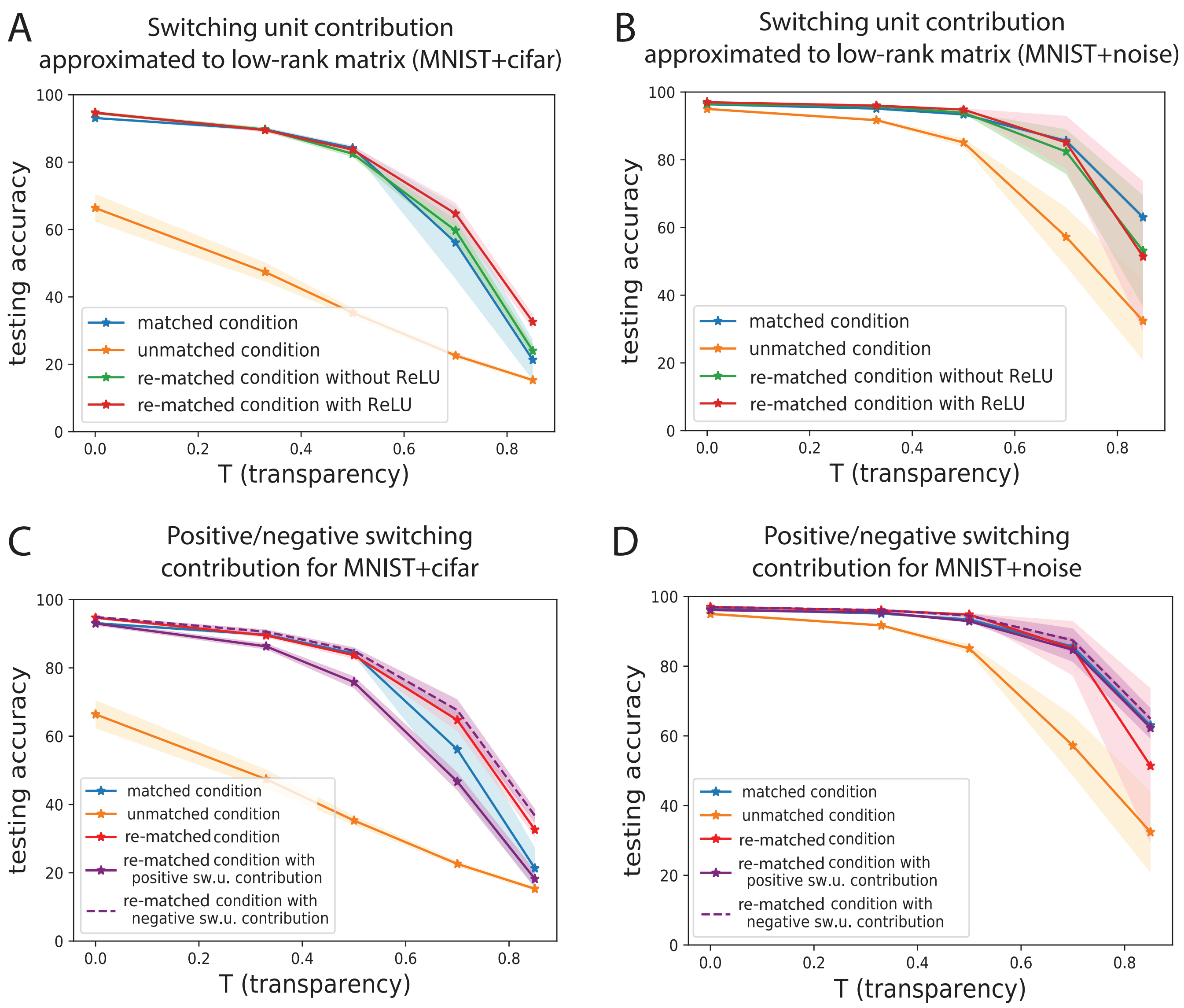

**Figure 8:** A) A switching network without using the ReLU nonlinearity for the switching unit activations on the MNIST+cifar dataset (green line) performs comparably well to the matched condition (blue line) and to the switching network with ReLUs described in main text Eq. (1) (red line). Without the ReLU non-linearity for the switching units, their contribution is low rank. Note that the basic network still has nonlinearities, they are removed here only from the switching units. B). Same as in A), but for MNIST+noise. C). Enforcing a strictly positive switching unit contribution for MNIST+cifar (solid purple line) decreases performance more than enforcing a strictly negative contribution (dashed purple line), p-value$< 2.23 \cdot 10^{-6}$ using the t-test. D). Same as C), but for MNIST+noise.

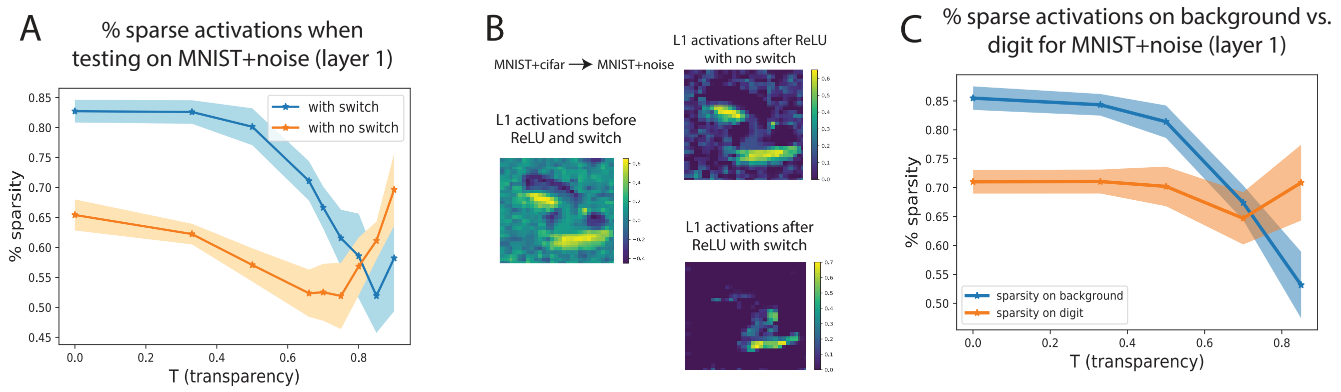

**Figure 9:** A). Sparsity (percentage layer 1 activations which are zero) when switching to MNIST+noise, with or without adding the switching contributions (blue vs orange line), and after applying ReLU. The mechanism described in Section 3.6 of the main text does not seem to apply for high transparencies when switching to MNIST+noise (sparsity % with switch is lower than the sparsity % without switch). B). Examples of (convolutional) layer 1 activation patterns with or without the switching contribution for $T = 0$, when switching to MNIST+noise. Figure 5B-C has shown these results for switching to MNIST+cifar. C). Sparsity (percentage layer 1 activations which are 0) within the background (blue) and within the digit (orange) when switching to MNIST+noise after adding the switching contributions and the ReLU non-linearity.

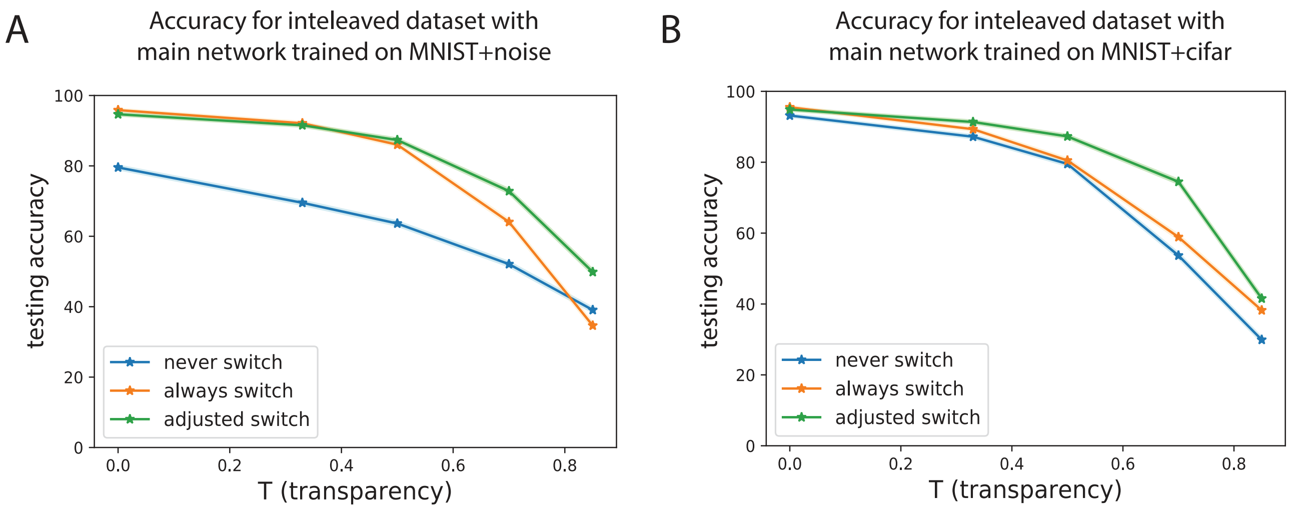

**Figure 10:** Accuracies for a network with a built-in "context detector", a simple one layer classifier trained on the data and used to decide whether the digit is set on noisy or CIFAR-10 context. This classifier outputs 0 for the first context (dataset 1) and 1 for the second context (dataset 2). The data to test this network contains interleaved MNIST+noise and MNIST+cifar images. A). Accuracy using the network with a built-in context detector, with the main network trained on MNIST+noise and switching units trained on MNIST+cifar. B). Same as A), but the main network is trained on MNIST+cifar and the switching units are trained on MNIST+noise.

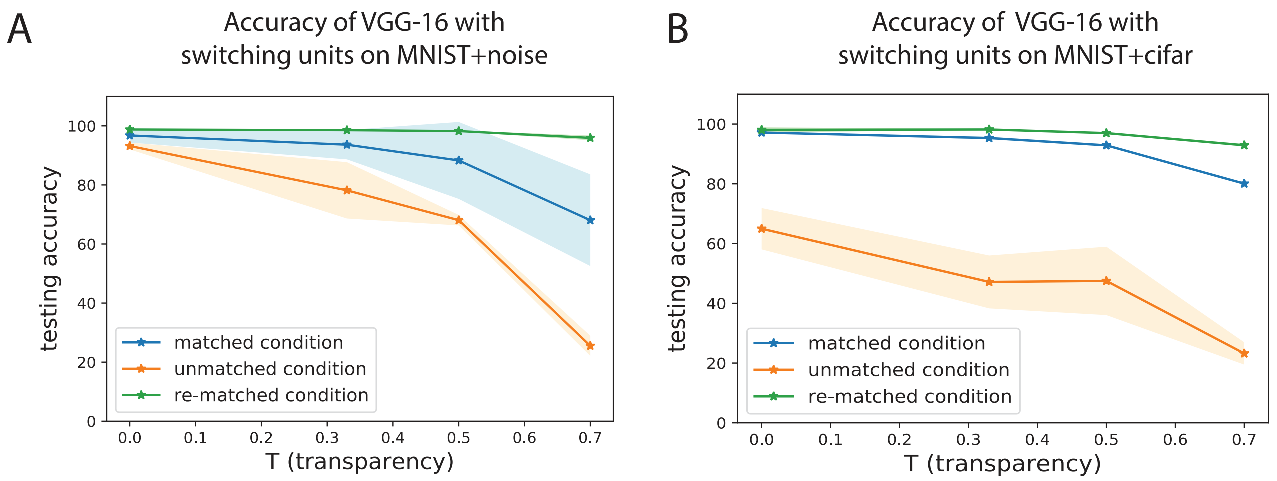

**Figure 11:** Switching units added to the VGG-16 deep network are effective for sequential context switching. A). Accuracy for the VGG-16 on MNIST+noise with switching units for the *re-matched condition* is superior to the accuracy for the *matched* and *unmatched conditions*. B). Same as in A), on MNIST+cifar.
